# Salience network expansion as a transdiagnostic neural marker for psychopathology risk in adolescence

**DOI:** 10.64898/2026.10.28.762645

**Authors:** Sanju Koirala, Robert Hermosillo, Kate J. Godfrey, Michael Anderson, Jacob Lundquist, Julia Moser, Eric Feczko, Oscar Miranda Dominguez, Kimberly B. Weldon, Isabella Linder, Maryam Mahmoudi, Lucina Q. Uddin, Steven M. Nelson, Jed T. Elison, Damien Fair, Brenden Tervo-Clemmens

## Abstract

Increased cortical surface area of the brain’s salience network (SAL) has been observed in depression and appears to emerge during early adolescence. However, similar expansion has also been reported in bipolar disorder (BD), obsessive compulsive disorder (OCD), and other atypicalities of salience network function have been broadly implicated across multiple psychiatric conditions. Whether increased SAL surface area is specific to depression or related more generally with psychopathology remains unclear. Using data collected as part of the Adolescent Brain Cognitive Development (ABCD) Study (baseline: N = 7942; ages 9–10; Year 2: N = 5901; ages 11–12), we quantified individual SAL size and examined associations with dimensional symptoms from the Child Behavior Checklist. SAL surface area showed substantial inter-individual variability, with expansion occurring primarily through encroachment into the default mode (DMN), action-mode (AMN), and frontoparietal (FP) networks. Larger SAL size showed a modest positive association with depressive symptoms in the baseline sample, consistent with prior reports; however, this association did not replicate in the year 2 internal replication sample. Instead, associations with total problems were stronger and replicated across both timepoints, suggesting a subtle, dimensional relationship between SAL organization and general psychopathology burden. These findings suggest that individual differences in SAL organization may represent a broader transdiagnostic neural correlate of dimensional psychopathology rather than a disorder-specific neural signature.

## Introduction

Increasing evidence links psychiatric disorders to alterations in large-scale functional brain network organization, with the salience network (SAL) consistently emerging as a key system of interest. Lynch and colleagues (2024) reported that SAL-associated frontal cortical surface area was approximately two-fold larger in individuals with major depressive disorder compared with controls, with similar expansion observed in bipolar disorder (Lynch et al. 2024). Complementing these findings, atypical SAL topography has been documented in treatment-resistant depression (Nahas et al.), while recent work reports comparable SAL expansion in individuals with obsessive-compulsive disorder (OCD) (Vaghi et al. 2025). Beyond topographic organization, alterations in SAL connectivity have also been implicated across a range of psychopathologies (Uddin 2014; Marshall et al. 2020; Green et al. 2016; Schimmelpfennig et al. 2023; Pannekoek et al. 2013; Willinger et al. 2024; Geng et al. 2015). Nevertheless, the relationship between individual differences in SAL topography and the dimensional structure of psychopathology remains poorly understood.

The SAL is a large-scale system involving the anterior insula (AI), dorsal anterior cingulate cortex (dACC), and prefrontal regions, including the inferior frontal and dorsolateral prefrontal cortices, as well as subcortical and limbic structures (Seeley et al. 2007). It is involved in detecting behaviorally relevant information and coordinating responses across cognitive, affective, and autonomic processes (Seeley et al. 2007; Menon and Uddin 2010; Uddin 2014). Through its interactions with systems supporting emotion, cognition, and internally directed processing, the SAL contributes to the integration of information that guides adaptive behavior (Molnar-Szakacs and Uddin 2022). These functions provide a rationale for examining SAL cortical surface area in relation to transdiagnostic dimensional measures of psychopathology, which capture symptom covariation across conventional diagnostic boundaries. However, whether individual differences in SAL size are associated with general transdiagnostic psychopathological burden or show specific relationships with particular symptom dimensions remains unknown.

Large-scale population neuroimaging provides a powerful approach to address this question. These samples capture broad variability in symptom severity and co-occurrence while providing the sample size and statistical power needed to examine associations between SAL organization and both individual symptom dimensions (e.g., depression) and broader psychopathology. This population-level dimensional approach directly complements precision fMRI studies (Gell et al. 2024), which have provided important insights into individualized network topography in deeply phenotyped clinical populations but often rely on diagnostic case–control comparisons. Such precision designs within individual diagnostic categories can be less suited to evaluating transdiagnostic patterns because diagnostic categories partition individuals according to clinical boundaries and may restrict the range of symptom severity and comorbidity represented within a sample (Cuthbert and Insel 2013). In contrast, population-based samples allow brain organization to be examined across continuously distributed dimensions of psychopathology, consistent with the Hierarchical Taxonomy of Psychopathology (HiTOP) (Kotov et al. 2017) and Research Domain Criteria (RDoC) (Cuthbert and Insel 2013; Morris et al. 2022) approaches. Thus, population-based neuroimaging provides an opportunity to determine whether individual differences in SAL topography are preferentially related to depressive symptoms or instead reflect broader variation in psychopathology.

Here, we leverage the Adolescent Brain Cognitive Development (ABCD) Study, a large, population-based developmental neuroimaging study, to examine whether individual differences in SAL size are preferentially associated with depressive symptoms or instead track multiple dimensions of psychopathology and general psychopathological burden. Using individualized cortical surface area estimates of canonical functional brain networks, we first examined associations between SAL size and Child Behavior Checklist (CBCL) symptom dimensions in the ABCD baseline sample (*N* = 7,942, ages 9–10). Given that brain–behavior associations are typically small, we used effect sizes from the baseline analysis to estimate the sample sizes required to detect effects of comparable magnitude in new samples, providing benchmarks for interpreting replication findings. We then evaluated the robustness and generalizability of the baseline associations using ABCD Year 2 assessment (*N* = 5,901, ages 11–12) as an internal replication sample and an independent, external replication sample from the Healthy Brain Network (HBN; *N* = 422, ages 5–21). Based on prior evidence of SAL expansion across psychiatric disorders and the broad functional roles of the SAL, we hypothesized that larger SAL size would be associated with greater symptoms across multiple domains of psychopathology and with general psychopathological burden, rather than showing a depression-specific association. Collectively, this study tests whether individual differences in SAL topography represent a neural correlate of dimensional, transdiagnostic psychopathology

## Results

### Salience network size exhibits substantial population-level variability, while its association with depressive symptoms is inconsistent

Salience network size showed substantial variability across individuals in both the baseline sample (mean = 11,270.62 mm^2^ [5.43%], SD = 4,511.56 mm^2^ [2.15%], range = 0–45,847.99 mm^2^ [0–22.66%]) and the year 2 internal replication sample (mean = 10,635.64 mm^2^ [5.14%], SD = 4,709.32 mm^2^ [2.24%], range = 0–39,851.30 mm^2^ [0–18.37%]). Proportional SAL size showed relatively modest reliability across the two-year interval (ICC [3,1] = 0.34, 95% CI = 0.31–0.36; Supplementary Figure 1), indicating partial temporal consistency in individual differences in SAL topography despite the expected within-person variability associated with estimating functional network boundaries from relatively short (10-minute) resting-state scans (Hermosillo et al. 2024).

Prior work reported larger SAL networks in individuals with major depression (Lynch et al. 2024). Consistent with this observation, larger SAL size was modestly associated with higher DSM-oriented depressive symptom scores in the baseline sample (*padj* = 0.040, ΔR^2^ = 0.00125; Figure 1B). However, this association did not replicate in either the year 2 internal replication sample or the external replication sample (both *padj* > 0.05; Supplementary Figure 4).

**Figure 1.**
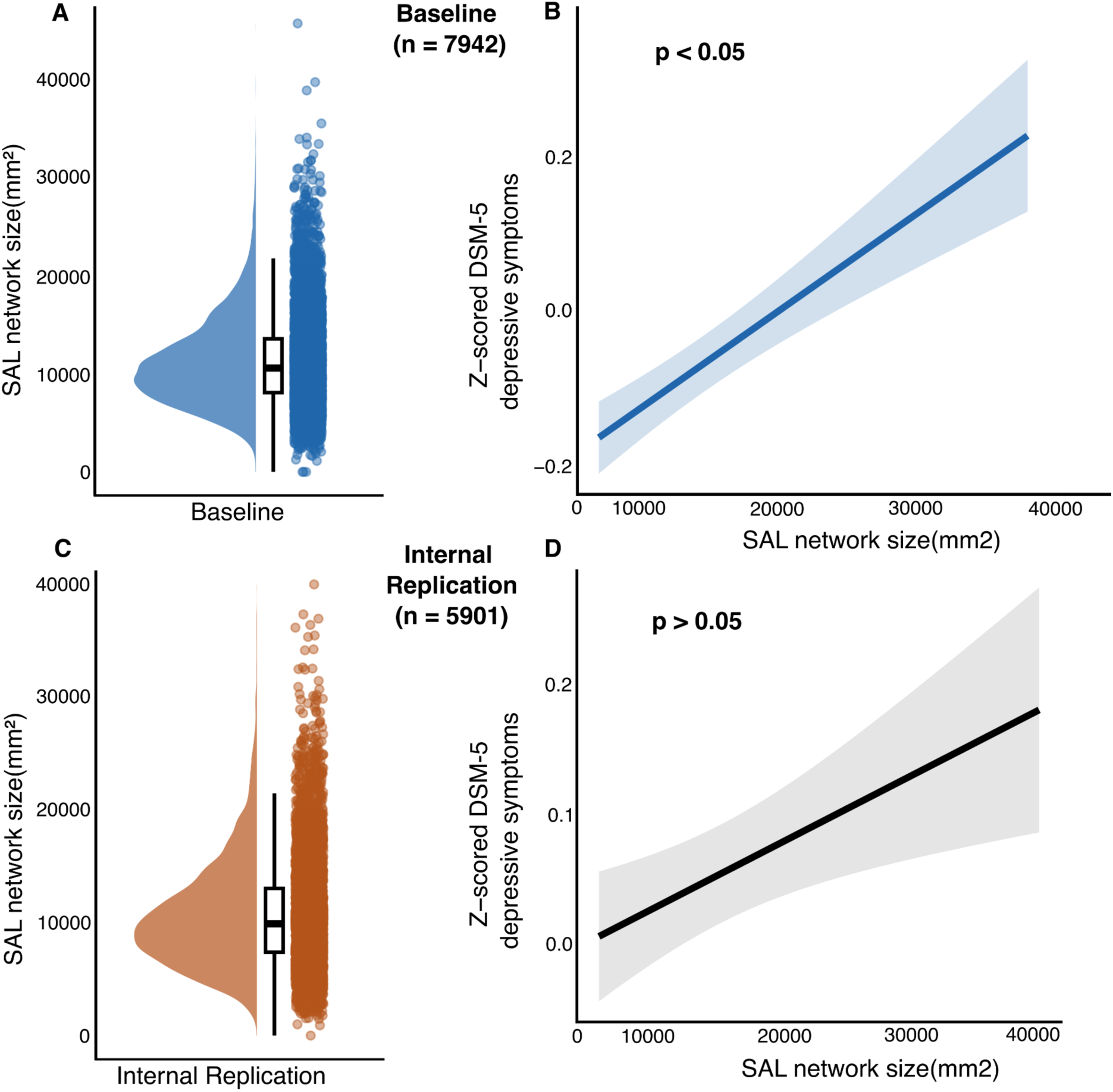
The association between salience network size and depressive symptoms is inconsistent. (A, C) Salience network size varies considerably across individuals in the baseline and internal replication samples, highlighting substantial heterogeneity in salience network topography during adolescence. (B, D) Symptom trajectories predicted by SAL size using GAMMs, standardized and offset for clarity. Larger SAL size was associated with higher depressive symptom scores in the baseline sample (B), but this association was not significant in the year 2 internal replication sample (D).

### Salience size relates to multiple symptom dimensions of psychopathology

We next examined associations between SAL size and a broader set of transdiagnostic symptoms. In the baseline sample, larger SAL size was significantly associated with higher social, attention, thought, and rule-breaking problems after multiple comparison correction, with modest effect sizes (all *padj* < 0.05; ΔR^2^ = 0.00058–0.00151). Associations with externalizing problems and DSM-oriented ADHD symptoms were positive but did not survive multiple comparison correction (*padj* = 0.0687 and 0.0622, respectively; Figure 1C, D).

In the year 2 internal replication sample, the overall pattern was similar, with positive associations across multiple symptoms. Associations with social problems, attention problems, externalizing problems, and DSM-oriented ADHD symptoms were significant (all *padj* < 0.05; ΔR^2^ = 0.00121– 0.00321). Other symptoms showed weaker associations that did not survive correction. SAL size was not significantly associated with DSM-oriented depression, anxiety, or OCD symptoms (all *padj* > 0.05; Figure 1E, F).

Overall, SAL size showed a broad pattern of small positive associations spanning internalizing, externalizing, and neurodevelopmental symptom dimensions rather than a specific DSM-oriented diagnostic domain. The variance explained was small, consistent with the magnitude of brain– behavior associations typically observed in large developmental neuroimaging studies (Marek et al. 2022). Together, these findings suggest that individual differences in SAL size may represent a transdiagnostic marker of psychopathology vulnerability during adolescence.

### Salience network size positively relates to general psychopathology

Building on symptom-specific associations (Figure 2), we next examined whether salience network (SAL) size predicted general psychopathology, operationalized as the CBCL Total Problems score. SAL size was significantly associated with total problems scores after multiple comparison correction (Figure 2A). Larger SAL size was associated with greater general psychopathology burden in both baseline (*padj* = 0.00046, ΔR^2^ = 0.00012) and the year 2 internal replication samples (*padj* = 0.00236, ΔR^2^ = 0.00126). These associations were consistently positive across both ABCD time points (Figure 2B). Together, these findings demonstrate that SAL size captures a transdiagnostic dimension of psychopathology severity in adolescence. Although SAL size was associated with multiple specific symptoms, its most consistent relationship was observed with general psychopathology burden rather than a single symptom domain such as depression.

**Figure 2.**
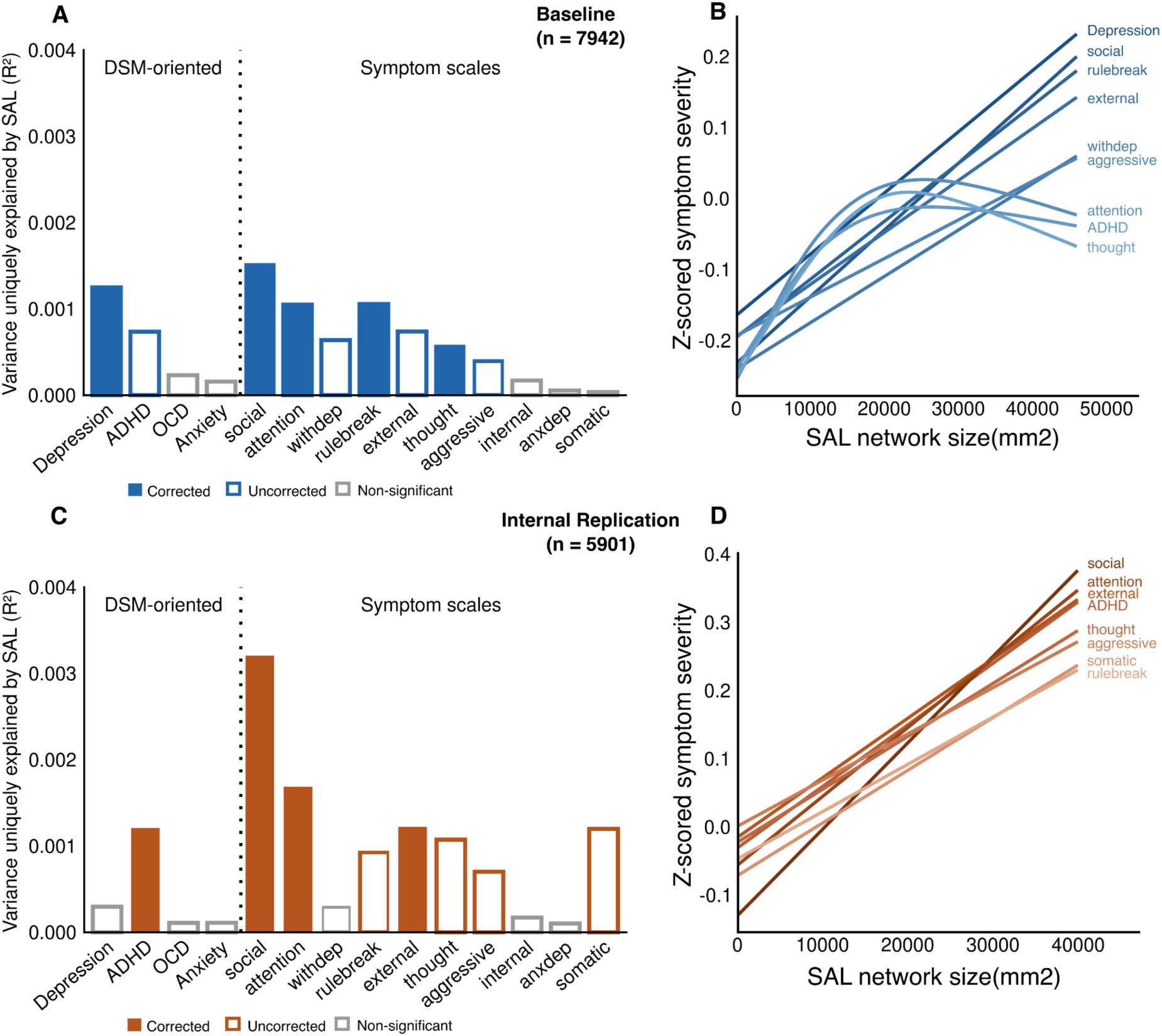
Variability in the size of salience network relates to multiple symptom specific dimensions of psychopathology. (A, C) Covariate-adjusted R^2^ values from GAMMS results indicate that variability in salience network size relates to multiple symptoms of psychopathology across baseline and internal replication samples. (B, D) Symptom trajectories predicted by SAL size using GAMMs for significant (p < .05) relationships, standardized and offset for clarity; larger SAL size is linked to higher symptom scores.

**Figure 3.**
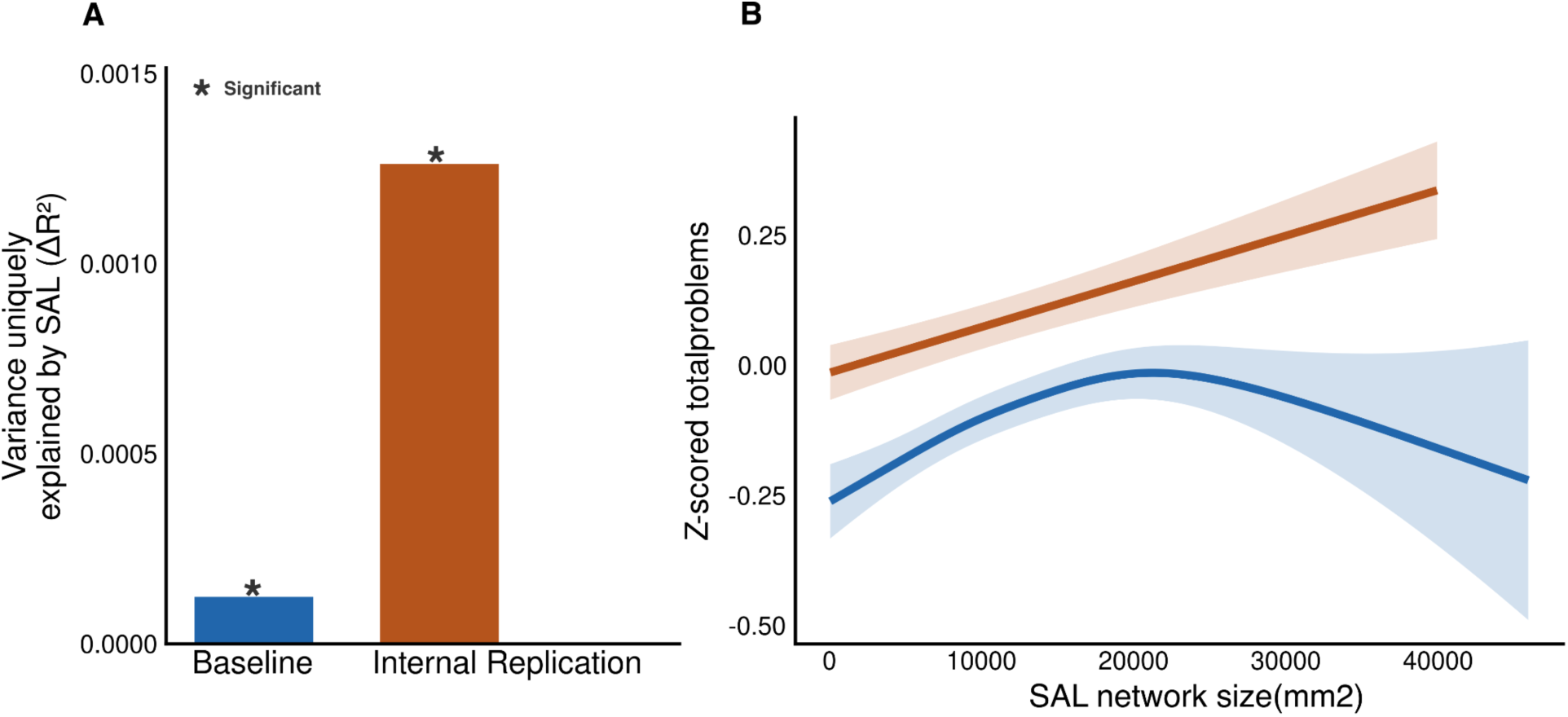
Salience network size predicts general psychopathology. (A) Covariate-adjusted R^2^ values from GAMMS results indicate that variability in salience network size predicts general psychopathology as indexed by CBCL total problem scores in both baseline and internal replication samples. (B) Standardized total problems trajectories predicted by SAL size using GAMMs, larger SAL size is linked to greater problem scores at both baseline and internal replication samples.

### Salience network is selectively enriched for positive psychopathology associations

Next, we tested whether the relationship between network size and psychopathology reflected a general property of cortical organization or was selectively concentrated in the salience network. Simply, does a larger SAL predict symptoms specifically, or does larger relative size of *any* network relate to psychopathology?

We examined the direction and consistency of associations between network size across a broad set of psychopathological symptoms for each of 15 canonical cortical networks (DMN, FP, etc). GAMMs predicting psychopathology symptoms from network size across major canonical cortical networks revealed that SAL exhibited the greatest enrichment for positive associations across both baseline and internal replication samples (Figure 4). Out of 15 total symptoms tested, SAL had significant positive associations with 5 of them at baseline and with 4 of them at internal replication samples, after adjusting for multiple comparisons.

**Figure 4.**
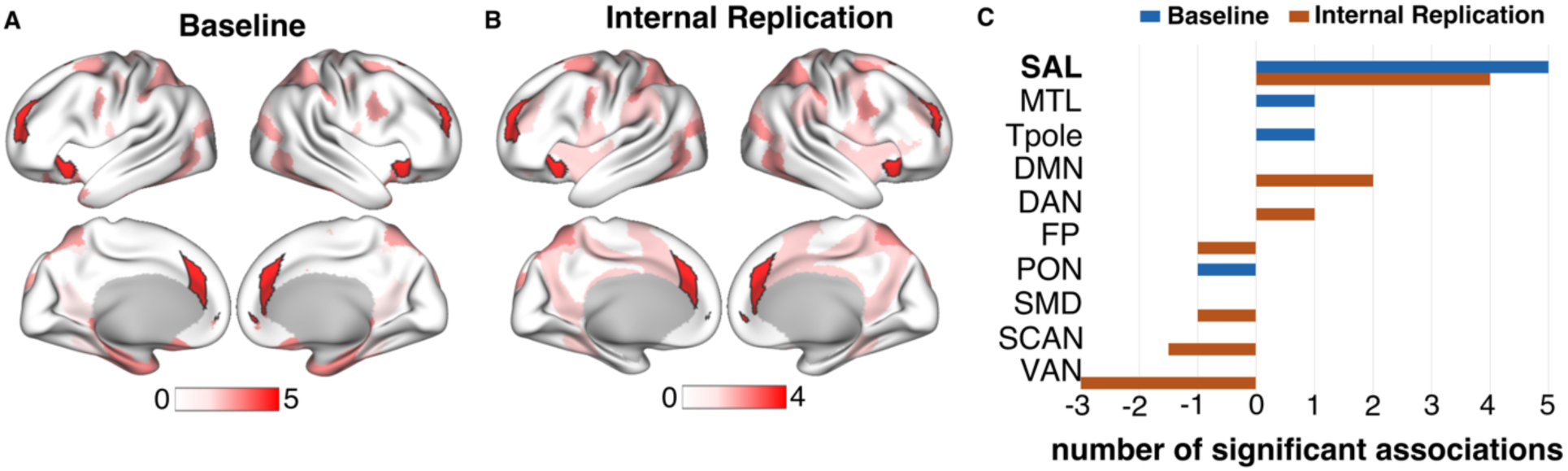
The salience network is enriched for positive psychopathology associations. (A, B) Network-level maps showing the number of psychopathology symptoms positively associated with network size. Outlined dark red patches denote SAL. Colors indicate the count of positive associations, with red denoting the highest number of positive associations; light pink and white indicate few or no significant effects. Across symptoms, SAL (parcels belonging to the SAL network outlined on the brain maps) demonstrates the greatest enrichment for positive associations relative to other networks. C) Number of significant associations across networks that showed any associations (positive or negative) with psychopathology symptoms.

**Figure 5.**
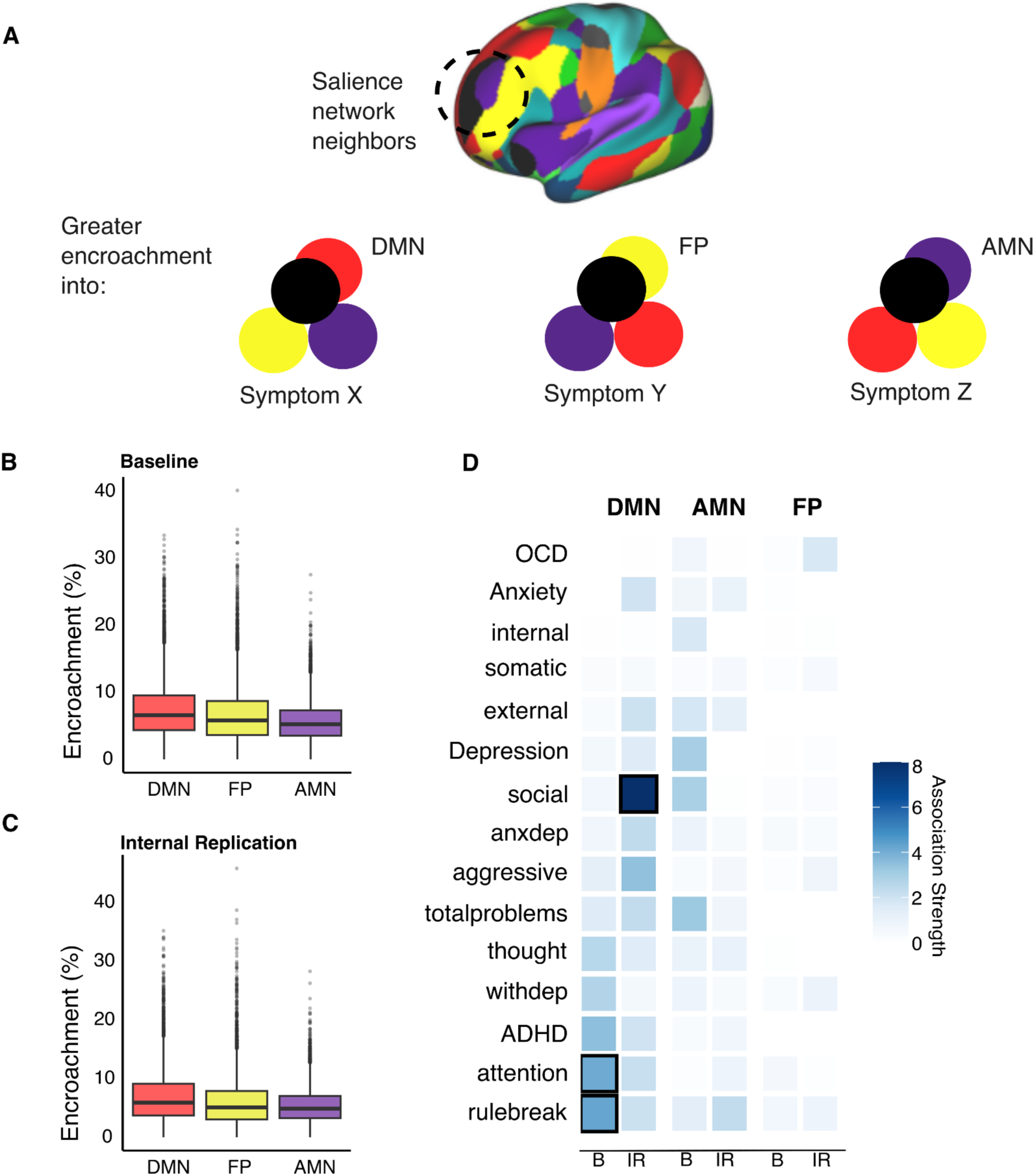
Patterns of SAL encroachment may shape symptom heterogeneity. *A)* Schematic illustrating the networks (FP, DMN, AMN) adjacent to SAL and potential encroachment patterns. Three hypothetical patterns illustrate how preferential encroachment into different neighboring networks may relate to distinct symptom profiles (X, Y, Z). *B, C)* Box plots showing the percentage of cortical area encroached upon by SAL within adjacent networks in the baseline (*B*) and internal replication (*C*) samples. *D)* Standardized *F*-values indexing the strength of associations from generalized additive mixed models (GAMMs), with SAL-encroached area within DMN, FP, and AMN entered as predictors of multiple psychopathology symptoms. Black-outlined squares indicate significant associations; unoutlined circles indicate non-significant associations. Overall, DMN encroachment showed the strongest and most consistent associations.

Compared to SAL, other networks showed fewer significant associations and were predominantly enriched for negative or null effects (Figure 4C). Importantly, this pattern was observed across multiple symptoms and in both samples, indicating that the positive associations were not driven by larger size in other networks or global cortical variation. Rather, variability in SAL size was specifically and consistently linked to elevated psychopathology across symptoms.

### Patterns of SAL encroachment may shape symptom heterogeneity

Prior work has shown that cortical expansion of the salience network (SAL) occurs through shifts in network boundaries, primarily encroaching on neighboring default mode (DMN), frontoparietal (FP), and action-mode (AMN) networks (Lynch et al. 2024; Vaghi et al. 2025). Consistent with these findings, SAL encroachment was observed across all three networks in both the baseline and internal replication samples (Fig. 4B–C). The distribution of encroachment differed across networks, with the greatest encroachment generally occurring into the DMN, followed by the FP and AMN networks. The full distribution of SAL encroachment across neighboring networks is provided in Supplementary Figure 3.

To examine whether the associations between SAL encroachment and psychopathology differed by target network, we used generalized additive mixed models (GAMMs) to estimate associations between encroached cortical area within the DMN, AMN, and FP networks and DSM-oriented and CBCL-derived symptom scores, controlling for demographic and imaging covariates. We used GAMMs F-statistics to characterize the strength of these semi-parametric associations.

As shown in Fig. 4D, SAL encroachment into the DMN showed the overall strongest associations with psychopathology at baseline. DMN encroachment was nominally associated with attention problems (*F* = 4.11, *p* = .04) and rule-breaking behavior (*F* = 4.25, *p* = .003), although these associations did not consistently survive correction for multiple comparisons. In contrast, associations involving AMN encroachment (*F* ≤ 3.2) as well as FP encroachment were uniformly weak (*F* < 1).

This pattern was broadly recapitulated in the year 2 internal replication sample, where DMN encroachment again showed the strongest magnitude and most widespread associations across symptoms. The strongest association was with social problems (*F* = 8.00, *p* = .004). Associations involving AMN encroachment remained generally weak, while FP encroachment showed limited associations, with OCD representing a comparatively notable exception (F’s < 2). Together, these findings suggest that although SAL encroaches on multiple neighboring networks, its associations with psychopathology are most pronounced for encroachment into the DMN.

## Discussion

### SAL topography captures a transdiagnostic dimension of psychopathology

Recent work has identified expanded salience network (SAL) topography in adults with depression and obsessive–compulsive disorder, raising the question of whether SAL expansion represents a disorder-specific or more general feature of psychopathology. In this large, population-based sample of adolescents, variability in SAL size showed inconsistent associations with depression but was more consistently related to dimensional measures spanning multiple symptoms of psychopathology. Notably, depression and OCD share features of repetitive negative and perseverative cognition, including rumination and difficulty disengaging from internally salient thoughts (Wahl et al. 2019; Moulds and McEvoy 2025; Weisholtz et al. 2017; Kaplan et al. 2018), suggesting that SAL variation may relate to processes that cut across diagnostic boundaries rather than to disorder-specific symptoms. Furthermore, associations were relatively strongest for total problems, suggesting that SAL topography may be more closely aligned with general psychopathological burden than with any single diagnostic category. Together, these findings extend prior adult work by suggesting that individual differences in SAL topography are evident during adolescence and may index a broad liability to psychopathology rather than a marker of a specific disorder. Acknowledging the modest magnitude of observed associations, a dimensional interpretation may therefore better capture the neurobiological relevance of SAL variation in adolescents than categorical diagnostic frameworks.

### SAL topography is enriched for psychopathology

The relative specificity of these associations to the SAL, compared with other large-scale cortical networks, further suggests that SAL topography may be particularly relevant to psychopathology. The SAL, anchored in the anterior insula and anterior cingulate cortex, integrates interoceptive, affective, and sensory information and coordinates interactions among systems supporting internally directed cognition and cognitive control (Menon and Uddin 2010; Seeley 2019). Individual differences in its cortical organization could therefore have broad connections to emotional regulation, attention, and cognitive control-processes implicated across diverse forms of psychopathology. Thus, rather than reflecting a disorder-specific vulnerability or sequelae, variation in SAL topography may represent a neural substrate through which multiple psychopathological processes are expressed. Future mechanistic work should determine the functional properties associated with individual differences in SAL size and organization.

### Patterns of SAL encroachment may shape symptom heterogeneity

SAL expansion was not spatially random but preferentially encroached upon specific canonical networks, most prominently the default mode (DMN), frontoparietal (FP), and action mode (AMN) networks, consistent with prior findings in adult clinical populations. These findings suggest that the functional implications of SAL expansion may depend not only on its overall size, but also on which networks are displaced or incorporated into its territory. In exploratory analyses, DMN encroachment showed the strongest and most consistent associations with psychopathology, whereas FP and AMN encroachment showed weaker or more symptom-specific relationships. Notably, although associations did not reach statistical significance, FP encroachment showed qualitatively larger association with OCD-related symptoms in the internal replication sample. This observation is of interest given prior findings of preferential FP encroachment in OCD (Vaghi et al. 2025), raising the possibility that different patterns of SAL encroachment may relate to distinct symptom profiles.

The triple-network model of psychopathology provides one potential framework for interpreting these findings (Menon 2011). This model emphasizes the roles of the SAL, DMN, and central executive network (CEN) in salience detection, internally directed cognition, and executive control, respectively, and proposes that altered organization and interactions among these systems may contribute to diverse psychiatric symptoms. In this context, the stronger associations observed for DMN encroachment raise the possibility that individual differences in the spatial organization of the SAL relative to systems supporting internally directed cognition may be particularly relevant to psychopathology. However, the present analyses do not directly assess functional communication between networks or the cognitive processes underlying these associations. Future studies integrating individual differences in network topography with functional connectivity and measures of specific psychological processes will be important for testing whether encroachment patterns have distinct functional implications.

These findings may pose questions for emerging circuit-based interventions that use neuroimaging-derived maps of network alterations to identify treatment targets for depression and bipolar disorder (Nahas et al.; Bambico et al. 2026; Siddiqi et al. 2024; Kong et al. 2025). If the spatial organization of the SAL varies meaningfully across individuals and different patterns of SAL encroachment are associated with distinct symptom profiles, investigators may consider whether group-level alteration maps fully capture the functional organization relevant to a given individual. Characterizing individual differences in SAL topography and their relationship to specific symptoms could therefore provide a foundation for more symptoms-informed approaches to circuit targeting, in which intervention sites are tailored to the functional organization and clinical profile of individual patients.

### Brain–behavior associations require large, well-powered samples

An important consideration is the modest magnitude of the observed brain–behavior associations. Although SAL size showed significant relationships with psychopathology in the ABCD cohort, the variance explained by these associations was small, consistent with contemporary large-scale studies of brain–behavior relationships (Marek et al. 2022; Tervo-Clemmens et al. 2023; Rosenblatt et al. 2023; Liu et al. 2023; Traut et al. 2022; Schulz et al. 2024; Winter and Depaoli 2022). Detecting such small effects reliably requires samples comprising several thousand participants (Supplemental Figure 5). Consistent with the presence of substantial individual variability in SAL organization, SAL size also varied considerably in the independent Healthy Brain Network (HBN) cohort. However, associations between SAL size and psychopathology were not significant in HBN, likely reflecting its substantially smaller sample size and potentially greater clinical, age and demographic heterogeneity (Supplementary Figure 5).

These findings underscore that individual differences in brain organization may be pervasive, while reproducibly linking these differences to behavior requires adequately powered samples. They also highlight a conceptual advantage of dimensional phenotyping: brain–symptom associations may more directly capture variation in specific psychological processes while avoiding heterogeneity introduced by grouping individuals with distinct symptom profiles into a single diagnostic category. This approach may therefore provide a more sensitive framework for identifying the functional relevance of individual differences in brain organization.

### Limitations and future directions

Several limitations should be considered. First, the cross-sectional nature of the primary analyses prevents longitudinal inference, and further studies are necessary to determine whether SAL expansion precedes symptom onset or reflects early manifestations of psychopathology. Longitudinal studies spanning adolescence into adulthood, including those leveraging forthcoming releases of the ABCD dataset, will be critical for determining whether SAL organization predicts subsequent clinical trajectories. Second, reliance on parent-reported measures may underrepresent certain psychopathology symptoms (Hope et al. 1999), highlighting the need for multi-informant and clinician-based assessments. Finally, although network size and encroachment capture spatial organization, they do not directly address fMRI temporal dynamics or functional interactions. Integrating measures of connectivity and network plasticity will therefore be important for determining the functional consequences of individual differences in SAL topography.

## Conclusion

Collectively, our findings suggest that SAL expansion represents a subtle, transdiagnostic feature of adolescent psychopathology. Rather than reflecting specific dimensional symptoms of depression or OCD, variability in SAL organization appears to associate best with general psychopathological burden in large population-based analyses and is accompanied by selective topographic redistribution into neighboring cortical systems, particularly the DMN. These findings identify large-scale cortical organization as a potential subtle marker of early psychiatric vulnerability and provide a framework for understanding how individual differences in brain topography may contribute to emerging mental health risk.

## Methods

### Participants and Datasets

We analyzed behavioral and resting-state fMRI data from the publicly available Adolescent Brain Cognitive Development Study (ABCD; release 6.0) and the Healthy Brain Network (HBN) datasets. After quality control and curation based on the availability of resting-state fMRI data and behavioral measures (see below), the final ABCD sample included N = 7,942 participants at baseline (ages 9–10 years) and N = 5,901 participants at Year 2 (ages 11–12 years). The HBN dataset included N = 422 participants ranging from 5 to 21 years of age.

### ABCD Study dataset

The ongoing ABCD Study® includes longitudinal annual behavioral and biennial fMRI data from 11,878 youth and their families across 21 sites in the United States (Casey et al. 2018). Study-wide exclusion criteria at the time of recruitment included standard MRI contraindications, history of traumatic brain injury, birth complications requiring more than 1-month hospitalization, a diagnosis of moderate to severe autism spectrum disorder, intellectual disability, major neurological disorders, schizophrenia, or a substance use disorder. Informed consent was obtained from a parent or guardian, and assent was collected from participants under the age of 18, in accordance with ethical standards and institutional review board (IRB) approval. More information about the ABCD study can be accessed through the official website: https://abcdstudy.org/.

We used the ABCD baseline visit (ages 9-10) for primary analyses. Owing to the relatively small effects we observed, we also tested the year 2 follow-up data as an estimate of inferential and model robustness. The initial ABCD sample included 11,878 participants. Analyses here used participants with at least 10 minutes of post-censored fMRI data with an FD threshold of < 0.2mm.

### HBN Dataset

For external replication and supplementary analyses, we used the Healthy Brain Network (HBN) data, which is a large transdiagnostic community sample, enriched for mental health concerns. HBN comprises over 3,000 children and young adults aged 5-21 years from four different sites. See Alexander et al. 2017 for additional detail.

### fMRI Acquisition

Acquisition parameters for the ABCD and HBN datasets are publicly available (Voldsbekk et al. 2023; Qu et al. 2025), and prior work suggests that these datasets provide sufficient cross-dataset comparability for research on individual differences. This project used resting-state fMRI data from the ABCD and HBN datasets. ABCD resting-state fMRI data were acquired with eyes open across multiple runs, with approximately 8 minutes per run and a target of approximately 20 minutes of total resting-state data per participant. HBN resting-state fMRI data were acquired with eyes open in two 5-minute runs, yielding approximately 10 minutes of resting-state data per participant.

### fMRI data processing

Details of acquisition for functional MRI data for the ABCD Study are described in (Casey et al. 2018). For processing, sequences were harmonized across Siemens, Philips and GE 3 Tesla (3T) scanners across all sites as outlined in Casey et al. 2018. ABCD fMRI data derivatives were obtained from the release 7.0 (https://docs.abcdstudy.org/latest/documentation/imaging/abcc_start_page.html#release-history) Analyses here use data from the ABCD-BIDS pipeline. ABCD-BIDS or (abcd-hcp-pipeline) is a BIDS App (Feczko et al. 2021) that adapts methods from the Human Connectome Project’s minimal preprocessing pipeline (Glasser et al. 2013). The DCAN Labs resting state fMRI analysis tools were used to output preprocessed MRI data in both volume and surface spaces. Brain extraction was performed by PreFreesurfer after denoising and bias field correction of the anatomical T1- and/or T2-weighted images. The DCAN-labs processing pipeline applies advanced normalization tools (ANTs) DenoiseImage to improve structural clarity and ANTs BiasFieldCorrection (ANTs) (Avants et al. 2009; Tustison et al. 2010) to reduce field bias(Marek et al. 2019). Further details can be found at https://docs.abcdstudy.org/latest/documentation/imaging/abcc_pipeline.html#abcd-bids-pipeline. The final ABCD analysis sample required participants with at least 10 minutes of low motion (FD < .2 mm) resting-state data, yielding a sample of 7942 participants for baseline and 5901 participants for year 2 internal replication sample.

HBN data was also processed using the ABCD-BIDS pipeline. Inclusion in the final analysis sample (N = 422) required participants to have a minimum of 8 minutes of low motion (FD < .2mm) resting-state data. We note a slightly shorter duration of time was required for inclusion in HBN, relative to ABCD, given that only 10 minutes of resting-state were collected.

### Template Matching: Individual Network Mapping

Following ABCD-BIDS preprocessing, individual-level functional network maps were generated using a template-matching approach following methods described in (Hermosillo et al. 2024). Briefly, resting-state fMRI data were motion-censored using a framewise displacement threshold of 0.2 mm, with outlier frames removed prior to connectivity estimation. Spatial smoothing was applied to the dtseries using a Gaussian kernel with σ = 2.55 mm. To ensure equivalent data duration across participants, frames were randomly selected to yield exactly 10 minutes of usable resting-state data per participant, including after removal of motion-related outlier frames.

For each participant, a whole-brain dense functional connectivity matrix (dconn) was generated from the resulting 10-minute motion-censored time series. The dconn was z-scored prior to template matching. Individual functional network organization was then estimated by comparing the spatial pattern of functional connectivity of each grayordinate with a set of canonical network templates using η^2^ (eta-squared). Each grayordinate was assigned to the network with which its connectivity profile showed the greatest similarity using a winner-take-all procedure.

The functional network templates were generated in prior work (Hermosillo et al., 2024) from ABCD Group 3 (ages 9–10 years), including 161 participants with 10 minutes of usable data. Before template matching, network templates were thresholded at Z > 1.0. The resulting template comprised 15 canonical functional networks: Default Mode Network (DMN), Dorsal Attention Network (DAN), Fronto-Parietal Network (FP), Visual Network (VIS), Action Mode Network (AMN), Somato-Cognitive Action Network (SCAN), Parietal-Occipital Network (PON), Parietal Memory Network (PMN), Salience Network (SAL), Ventral Attention Network (VAN), Medial Temporal Lobe Network (MTL), Temporal Pole Network (TPOLE), Auditory Network (AUD), Somato Motor Lateral Network (SML), and Somato Motor Dorsal Network (SMD).

Following winner-take-all assignment, small isolated network regions were cleaned by removing islands smaller than 30 grayordinates. For each removed island, the network assignment was replaced with the modal network assignment among its neighboring grayordinates, thereby incorporating the island into the surrounding network. The resulting individualized network maps were used for subsequent quantification of network topography and size.

### Network Surface Area

Functional network surface area was quantified as the cortical surface area occupied by each network. First, vertex-wise areas were first estimated on the individual midthickness surface using the workbench command: wb_command-surface-vertex-areas. Next, for each functional network, we summed the areas of all assigned vertices to obtain its surface area. The surface area values of all 15 networks were summed to create the total surface area of the cortex. This total cortical surface area was used in subsequent analyses as a covariate to account for inter-individual variability in total cortical surface area.

### Encroachment Calculation

We first generated a group-average functional network map using independent data from the Human Connectome Project–Development (HCP-D; ages 8–15, n = 121). For each individual in ABCD, we then identified vertices assigned to the salience network that did not spatially overlap with the salience network in the independent group-average map (Supplementary Figure 2). These voxels were defined as “encroached” voxels. Next, for each encroached vertex, we determined its canonical network assignment based on the group-average map. We then quantified the area of these vertices within each network, yielding a measure of “encroached area” per network for each individual.

### Behavioral Measures

#### CBCL Scores

Psychopathology symptom scores for both ABCD and HBN were obtained using the Child Behavior Checklist (CBCL; Achenbach System of Empirically Based Assessment, ASEBA) for both studies. The CBCL is a widely used, standardized parent-report questionnaire designed to assess a broad range of emotional, behavioral, and social problems in children and adolescents aged 6–18 years (with parallel forms for younger ages depending on dataset version).

The CBCL yields empirically derived scores across multiple levels of organization, including syndrome scales, DSM-oriented scales, and broadband composites. This analysis used age- and sex-normed T-scores available in the release, with higher scores indicating greater symptom severity. T-scores were used for the analyses in this work. The following syndrome scores as well as DSM-oriented scores were used for the ABCD study datasets. For HBN data, only these syndrome scales were used as DSM-oriented scales were not available.

Syndrome Scales: The CBCL syndrome scales reflect empirically derived dimensions of co-occurring behavioral and emotional problems which capture dimensional patterns of psychopathology rather than categorical diagnoses. These syndrome scales include Anxious/Depressed, Withdrawn/Depressed, Somatic Complaints, Social Problems, Thought Problems, Attention Problems, Rule-Breaking Behavior, and Aggressive Behavior.

DSM-Oriented Scales: In addition to syndrome scales, the CBCL provides DSM-oriented scales. These scales were developed through expert clinician ratings to align CBCL items with DSM diagnostic categories, which provide. These scales provide dimensional approximations of DSM-based symptoms. These include Depression scores, ADHD scores, OCD scores, and Anxiety scores. These scales provide dimensional approximations of DSM-based symptoms and are commonly used in developmental psychopathology research.

#### Covariates

All statistical models (described below) included a consistent set of covariates to account for potential confounding effects. In ABCD data, covariates included age, sex, socioeconomic status (SES), site of data collection, and family relatedness. Age reflected the participant’s developmental stage (approximately 9–10 years at baseline and 11–12 years at internal replication sample). Sex was coded as male or female. SES was operationalized using parental education. The site was included to account for variability across 21 data collection sites in the United States, and family ID was included to account for non-independence among siblings. For HBN data, covariates included age, sex, socioeconomic status, and site of data collection.

#### Analytic Strategy

All analyses were conducted using MATLAB 2019b (The MathWorks, Natick, MA) and RStudio version 2023.12.1+402. Generalized Additive Mixed Models (GAMMs) were used to examine associations between individual differences in salience network (SAL) size and dimensional psychopathology. Models included age, sex, and socioeconomic status (parental education) as fixed-effect covariates. To account for the non-independence of observations arising from the multisite and family-based structure of the ABCD Study, random intercepts were specified for site and family nested within site.

#### GAMMs: Modeling relationship between SAL size and psychopathology

Separate GAMMs were fitted for each symptom to estimate its fixed-effect association with SAL size, while accounting for covariates and random effects. GAMMs were selected to flexibly model potential nonlinear relationships between brain measures and age while accounting for hierarchical data structure. All models included SAL size as the primary predictor and symptom scores as outcome variables. Age, sex, and socioeconomic status (SES) were included as fixed-effect covariates. To account for non-independence in the data, random intercepts were included for family and site of data collection. All GAMMs were implemented using the default thin plate regression spline smooth terms (tp), allowing for flexible modeling of age-related effects without imposing a strict parametric form.

Model performance was quantified using changes in marginal R^2^ to estimate the variance in symptom scores uniquely explained by SAL size. Specifically, for each model, two GAMMs were fit: one including SAL size as a predictor and a reduced model excluding SAL size but retaining all covariates (age, sex, and socioeconomic status). The incremental variance explained by SAL was then calculated as the difference in marginal R^2^ between the full and reduced models (ΔR^2^ = R^2^_full − R^2^_reduced). This approach allowed for estimation of the unique contribution of SAL size beyond covariates and random effects. Results were corrected for multiple comparisons using FDR correction (corrected p < 0.05). To characterize the direction and functional form of associations, fixed-effect model-derived predictions were generated from fitted GAMMs. Psychopathology scores were z-scored within each dataset to facilitate comparability across symptoms and samples. Total problems scores were standardized across the combined baseline and Year 2 samples prior to model estimation. This transformation placed symptom severity on a common scale while preserving the relative ranking of individuals within the overall sample distribution, allowing comparison of model-derived trajectories across visits. Predictions were obtained across the observed range of SAL while holding covariates constant. Continuous covariates (age, SES, and brain size) were fixed at their sample means, and the categorical variable (sex) was fixed at its modal category. Random effects were held constant at a reference level during prediction. This procedure yielded predicted values representing the SAL–symptom association for a standardized reference individual. Predicted values and corresponding standard errors were extracted from the GAMM smooth terms and used to visualize the functional relationship between SAL size and symptom severity. These model-derived estimates were plotted as smooth curves with 95% confidence bands (±1 SE) for each dataset, allowing for comparison of effect direction, magnitude, and potential nonlinearity across symptoms and samples. All visualization and prediction procedures were implemented in R using the *mgcv* and *ggplot2* packages.

#### SAL Enrichment Summary

To determine whether brain–behavior associations were preferentially concentrated within specific large-scale functional networks, we conducted a network-level analysis across 15 canonical cortical networks and all tested psychopathology symptoms. For each network– symptom pair, GAMMs were fit using network surface area as the predictor and psychopathology as the outcome, with the same covariate and random-effects structure described above. Statistical significance was assessed for the network-size term, and *p*-values were corrected across all network–symptom tests using the Benjamini–Hochberg false discovery rate (FDR) procedure.

For each FDR-significant association, the direction of the network–psychopathology relationship was determined from the model-derived association across the observed range of network size. Because the fitted relationships were largely monotonic, associations were classified as positive or negative based on their corresponding linear best approximation. Network-level enrichment was then quantified as the number of FDR-significant positive and negative associations observed for each network. Enrichment was defined as a greater concentration of significant positive associations within a given network relative to the other networks.

#### Linking SAL Encroachment Patterns to Psychopathology

To examine associations between SAL encroachment onto other functional networks and psychopathology, DMN, FP, and AMN encroachment areas were included simultaneously as smooth predictors in separate models for each symptom, following the GAMM framework described above. Model inference was based on smooth-term F-statistics and associated *p*-values, with Bonferroni correction applied for multiple comparisons. The same analysis was performed separately in the baseline and internal replication samples. Results were visualized as heatmaps of F-statistics across encroachment networks and psychopathology symptoms, with annotations indicating significance.

## Supplementary Information

**Supplementary Figure 1.**
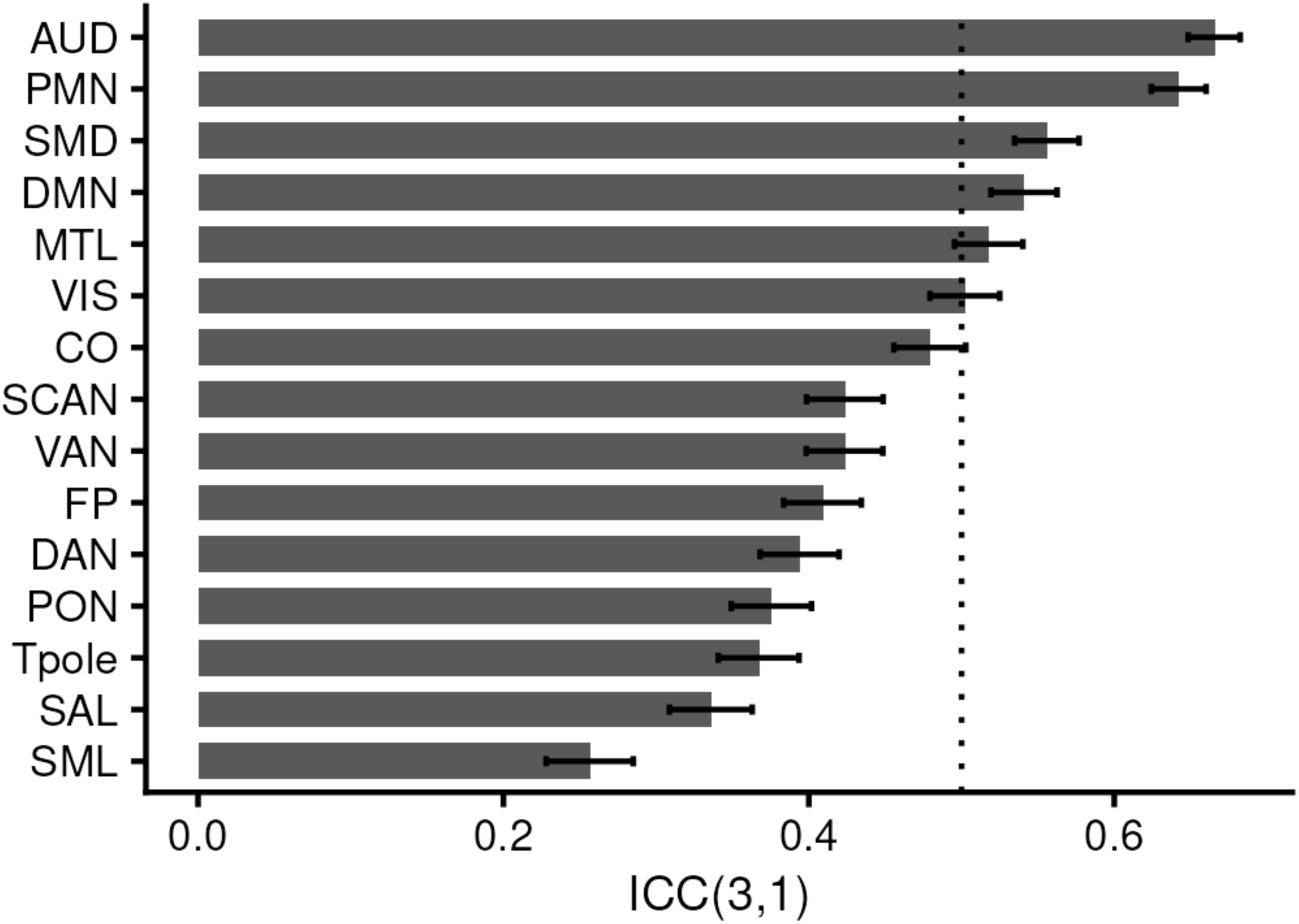
Longitudinal reliability of proportional functional network size. Intraclass correlation coefficients (ICC) were calculated for proportional network size (network size normalized by total brain network size) between baseline and Year 2 assessments among participants with available data at both time points. Bars represent ICC estimates, and error bars indicate 95% confidence intervals. The dotted horizontal line denotes an ICC value of 0.50, shown as a reference for moderate reliability

**Supplementary Figure 2.**
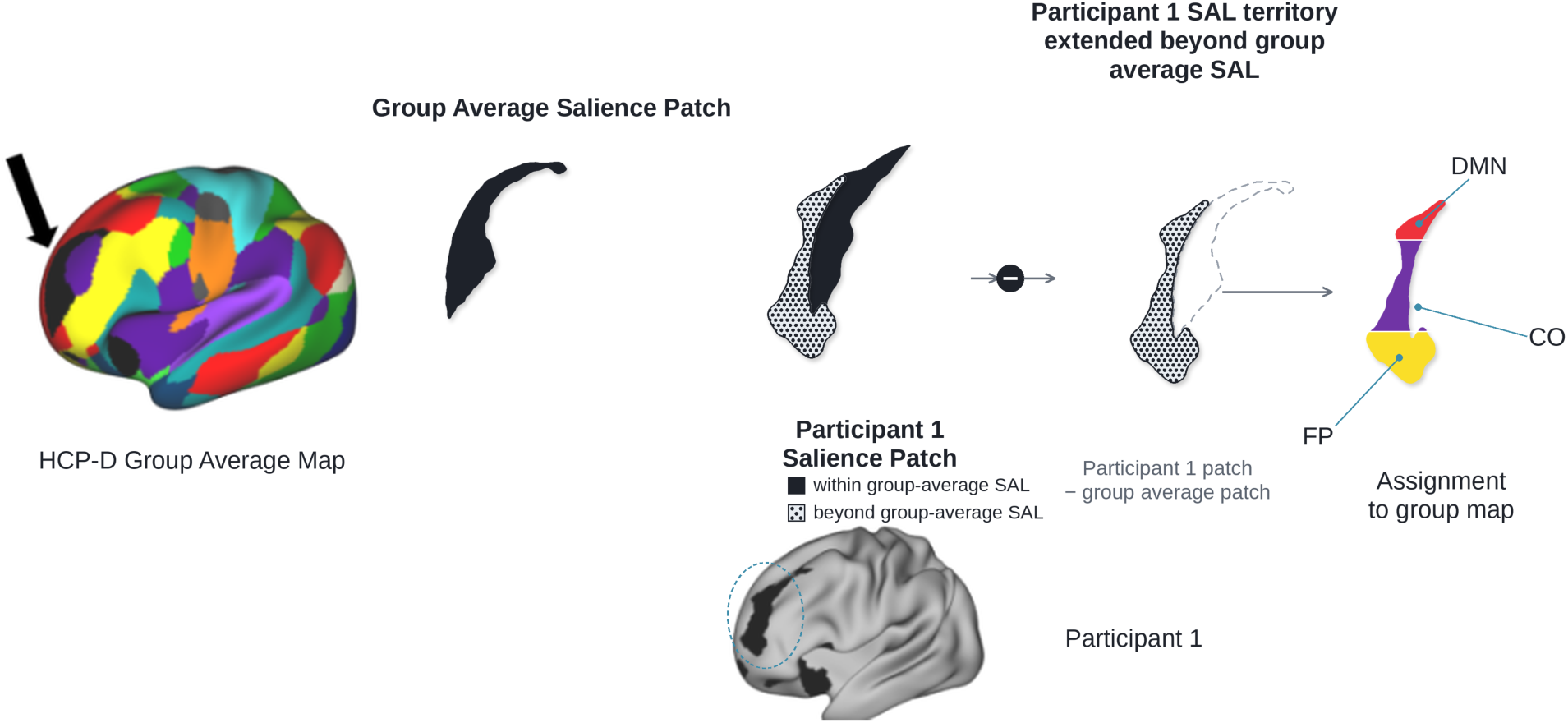
Encroachment Calculation. The group average map was generated using Human Connectome Project-Developmental (HCP-D). For each individual, we then identified voxels assigned to the salience network that did not spatially overlap with the salience network in the group-average map. These voxels in the participant’s SAL territory that extended beyond the group average SAL were defined as “encroached” voxels. For each encroached voxel, we determined its canonical network assignment based on the group-average map. We then quantified the area of these voxels within each network, yielding a measure of “encroached area” per network for each individual.

**Supplementary Figure 3.**
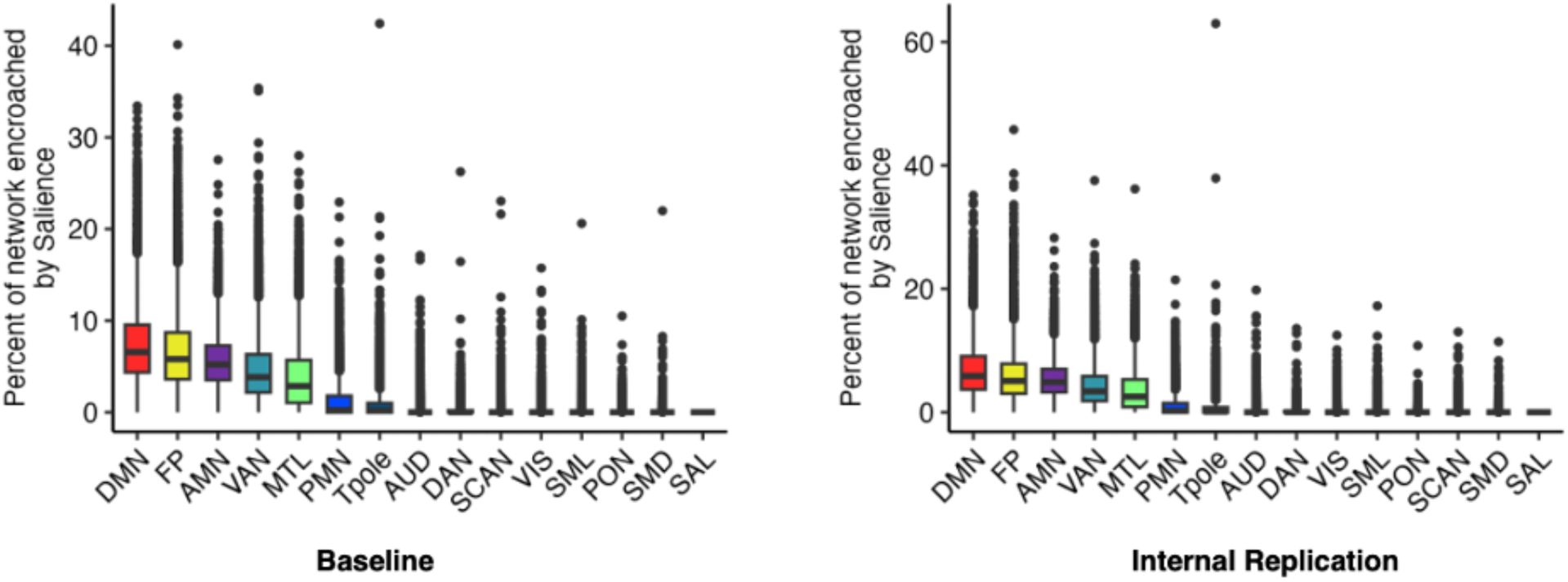
Distribution of cortical network area encroached upon by the salience network (SAL). Box plots showing the percentage of cortical area within each canonical functional network that is encroached upon by the SAL, shown separately for the baseline (left) and internal replication (right) samples. The distribution of SAL encroachment was greatest within the default mode network (DMN), frontoparietal network (FP), and action mode network (AMN) in both samples.

**Supplementary Figure 4.**
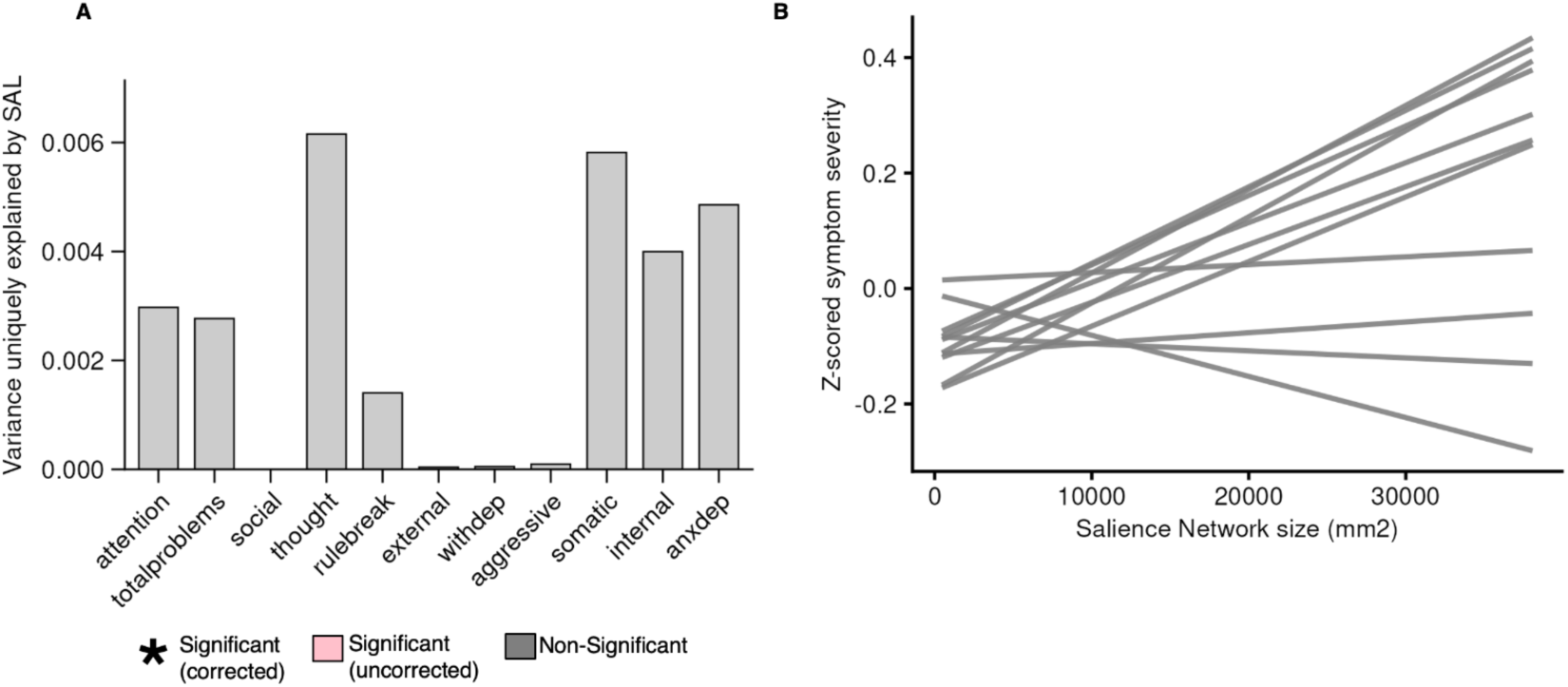
Relationship between Salience network and various symptom specific dimensions of psychopathology in Healthy Brain Network data. (A) Covariate-adjusted R^2^ values from GAMMS results indicate that variability in salience network size is not significantly associated with dimensions of psychopathology. (B) Symptom trajectories predicted by SAL size using GAMMs standardized and offset for clarity; despite non-significance, larger SAL size is linked to higher symptom scores.

**Supplementary Figure 5.**
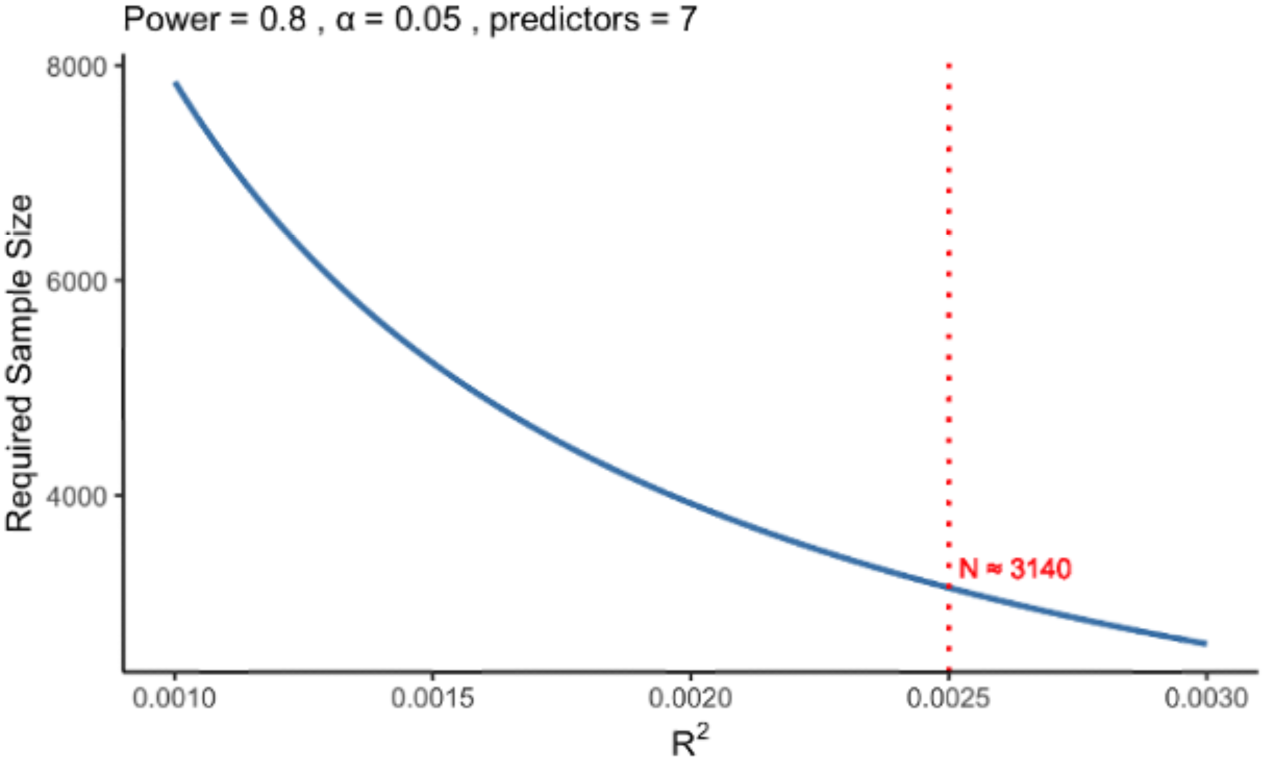
Sample size requirements for detecting small brain–behavior associations. Required sample size is plotted as a function of the proportion of variance in psychopathology explained by salience network (SAL) size (R^2). Sample size requirements were estimated assuming 80% statistical power, a two-sided significance level of alpha = 0.05, and seven predictors, corresponding to the predictors and covariates included in the GAMM framework. The analysis was evaluated across a range of small effect sizes (R^2 = 0.001–0.003) encompassing the effects observed in the present ABCD analyses. The vertical dashed line indicates a representative effect size of R^2 = 0.0025, corresponding to an estimated required sample size of approximately N = 3,140. This analysis illustrates the large sample sizes required to detect brain–behavior associations of the small magnitude observed in the present study and provides a basis for planning future replication studies.

## Acknowledgements

We are grateful to participating families and ABCD research staff for their time and effort.

## Data and Code Availability Statement

Data used in the preparation of this article were obtained from the Adolescent Brain Cognitive Development Study® (http://abcdstudy.org). Researchers with an approved NIH Brain Development Cohorts (NBDC) Data Use Certification (DUC) may obtain ABCD Study data. The analysis code will be publicly available upon publication.

